# Metacommunity in dynamic landscapes

**DOI:** 10.1101/021220

**Authors:** Charles Novaes de Santana, Jan Klecka, Gian M. Palamara, Carlos J. Melián

**Author notes:** **Author contributions:** CNdS and CJM designed the study and drafted the initial model. CNdS and JK implemented the model. All authors analyzed the data and contributed substantially to initial draft and revisions of the manuscript. Shared first authors. Corresponding author: Charles Novaes de Santana. University of Zurich, Institute of Evolutionary Biology and Environmental Sciences, Building Y27-J-54, Winterthurerstrasse 190, CH-8057 Zürich, Switzerland.

## Abstract

Predictions from theory, field data, and experiments have shown that high landscape connectivity promotes higher species richness than low connectivity. However, examples demonstrating high diversity in low connected landscapes also exist. Here we describe the many factors that drive landscape connectivity at different spatiotemporal scales by varying the amplitude and frequency of changes in the dispersal radius of spatial networks. We found that the fluctuations of landscape connectivity support metacommunities with higher species richness than static landscapes. Our results also show a dispersal radius threshold below which species richness drops dramatically in static landscapes. Such a threshold is not observed in dynamic landscapes for a broad range of amplitude and frequency values determining landscape connectivity. We conclude that merging amplitude and frequency as drivers of landscape connectivity together with patch dynamics into metacommunity theory can provide new testable predictions about species diversity in rapidly changing landscapes.

## Introduction

Metacommunity theory provides a number of insights into the role of dispersal for species coexistence in landscapes composed of units of suitable and unsuitable habitats (Holyoak *et al.* (2005)). Empirical studies have largely focused on dispersal rates with only recent emphasis on patterns of landscape connectivity (Kneitel & Chase (2004); Cadotte (2006)). Most studies have shown that increasing connectivity tends to increase persistence and richness (Ellner *et al.* (2001); Fox *et al.* (2011)), but examples of decreasing richness with increasing connectivity are also known (Davies *et al.* (2009); Altermatt *et al.* (2011)). Theoretical models predict that habitat loss and fragmentation may reach a threshold beyond which there is a rapid avalanche of species extinctions (Fahrig (2002); Ovaskainen & Hanski (2003); Rybicki & Hanski (2013)). These predictions gained empirical support from studies of deforestation where a transition from a continuous forest to more isolated and smaller fragments of the original habitat occurs and is accompanied by significant species loss (Laurance *et al.* (1997); Metzger *et al.* (2009)). Fluctuations in landscape availability (random or seasonal) are also common in nature (Sprugel (1991); Ruiz *et al.* (2014)) but the consequences of fluctuations in landscape connectivity for species richness received less attention, with the exception of disturbances (Sousa (1984); Supp & Ernest (2014)). Whether landscape connectivity increases or decreases persistence and regional species richness, dispersal abilities of organisms, which define habitat connectivity, are affected by the fluctuations in the environment and various habitat characteristics. Many of these factors fluctuate with different frequencies, with some showing high intraday variation while others fluctuate daily, seasonally or at larger time scales (Stenseth *et al.* (2002)).

Landscape dynamics encompasses two major processes: patch dynamics and variation in landscape connectivity. Patch dynamics is defined as changes of the number and position of patches, changes of patch habitat characteristics, size and suitability. Fluctuations in landscape connectivity are defined as changes of the matrix organisms have to cross to disperse from one patch or habitat to another (figure 1 and table 1 for a glossary of the main concepts). In some cases, temporal and spatial dynamics are correlated. Examples include fire size distributions with highly frequent small-scale fires and rare large-scale ones (Hantson *et al.* (2015)). At large temporal scales, transitions between habitat types at the continental scale occur during glacial-interglacial cycles (Werneck *et al.* (2011)). There are also examples of large-scale landscape dynamics over short time scales, such as daily tides and seasonal changes of sea ice extent (see animations SI-A1 and SI-A2). Correlated and uncorrelated temporal and spatial scales driving landscape dynamics may have implications for metacommunity dynamics. For example, landscape dynamics in combination to climate change velocity may impact threatened populations (Loarie *et al.* (2009)), affect population divergence and speciation (Aguilée *et al.* (2011)), and drive changes of the latitudinal biodiversity gradient over time (Mannion *et al.* (2014)).

**Table 1.**
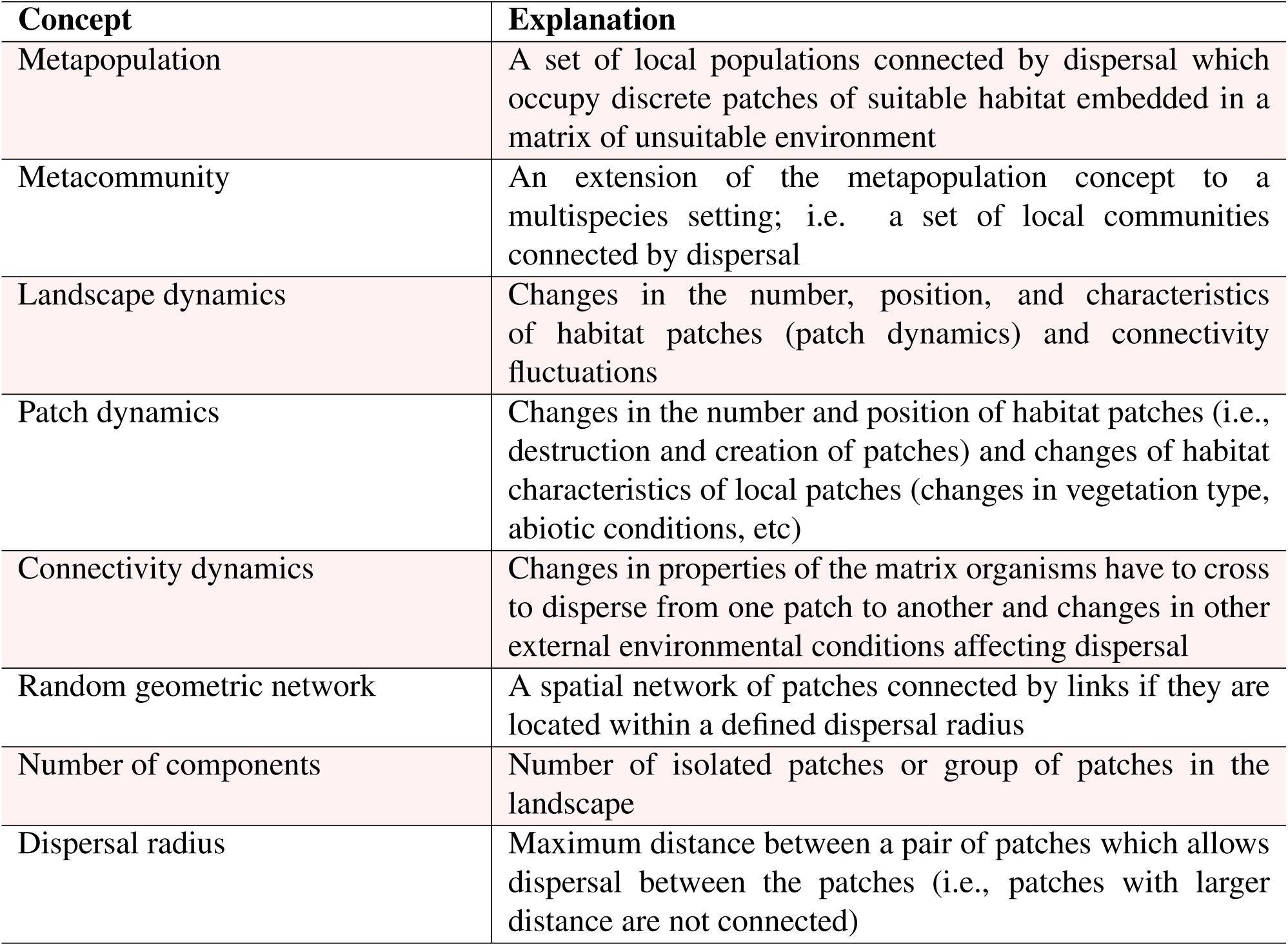
Glossary of concepts

**Figure 1:**
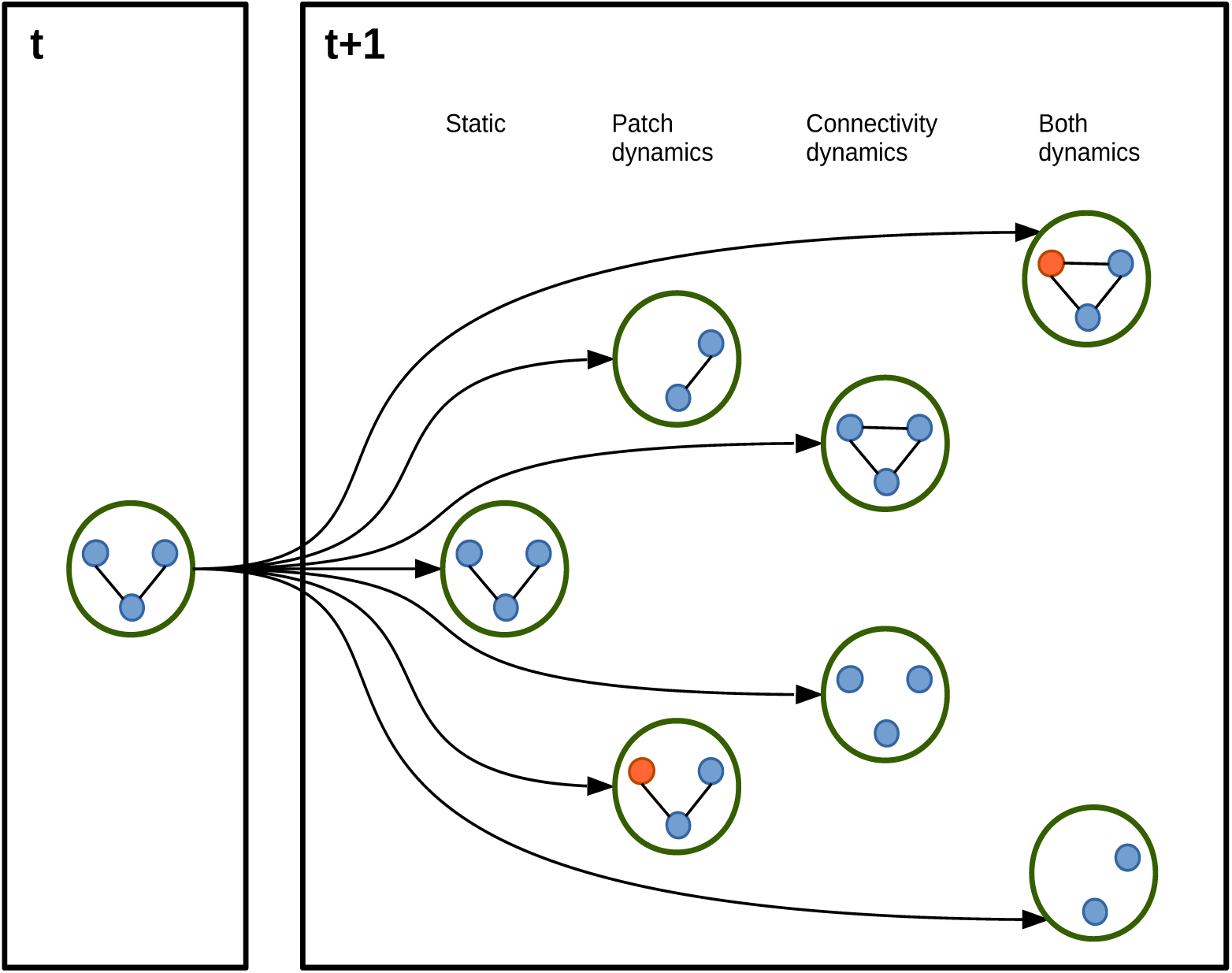
Two major processes of landscape dynamics: Patch dynamics represents changes in the number and position of patches, changes of patch habitat characteristics, size and suitability. Connectivity dynamics represents changes in the landscape matrix in future time points. Both dynamics, patch and connectivity dynamics, may happen at the same time.

Patch dynamics, i.e. the process of destruction of patches and appearance of new ones, has been addressed by numerous theoretical studies of metapopulations (Hanski (1999); Cornell & Ovaskainen (2008); Drechsler & Johst (2010)). Hanski (1999) derived formulas for predicting patch occupancy of a single population in landscapes characterized by temporal patch dynamics. The mean species lifetime in a network of dynamical patches can also be estimated (Drechsler & Johst (2010)). Recent studies have shown that the rate of patch turnover is critical for metapopulation persistence. For example, Reigada *et al.* (2015) showed that increasing the rate of patch dynamics decreases metapopulation persitence when dispersal is continuous, while persistence is facilitated by pulsed dispersal. The links connecting different patches can also vary in time. For example, the connectivity of habitat patches in the polar regions fluctuates seasonally according to sea ice extent (see animations SI-A1 and SI-A2). Connectivity dynamics can therefore be critical in determining landscape structure. However, connectivity dynamics has received less attention in metacommunity and metapopulation ecology (Holyoak *et al.* (2005); Johst *et al.* (2011)). The concept of connectivity dynamics has been more commonly used in disease ecology (Dushoff *et al.* (2008); Keeling & Eames (2008); Ross (2010)). For example, sinusoidal forcing of the transmission rate can accurately describe fluctuations of incidence in the host observed in the dynamics of the host–influenza system (Dushoff *et al.* (2008)).

Despite the scarcity of theoretical predictions, there is empirical evidence that connectivity dynamics may play an important role for dynamics of metapopulations in heterogeneous landscapes. Most of the empirical evidence comes from studies which focused on single-species metapopulation persistence where habitat connectivity is driven by the characteristics of the landscape matrix separating habitat patches as perceived by the organisms (Eycott *et al.* (2012)). For example, dispersal of amphibians between ponds is strongly affected by the terrestrial habitat separating the ponds (Buskirk (2012); Cline & Hunter (2014)) and by weather (e.g., moisture) (Rittenhouse *et al.* (2009)). Similarly, dispersal of butterflies also depends on the landscape matrix (Kuefler *et al.* (2010)) and dispersal kernels fluctuate in time (Schtickzelle *et al.* (2012)). In fish, interconnections between rivers forming during periods of heavy rain can connect otherwise disconnected habitats and allow for dispersal and gene flow (Boizard *et al.* (2009)). Here we connect temporal and spatial changes of landscape connectivity to metacommunity dynamics and species richness. We use amplitude and frequency as a proxy to describe both spatial and temporal fluctuations in the landscape, varying periodically the dispersal radius of the organisms (i.e., any two patches are connected if their distance is lower or equal than the dispersal radius, figure 2 and table 2 for the parameters used). We then compare landscapes with no connectivity change (i.e., static landscapes) with landscapes whose dispersal radius fluctuates with a given amplitude and frequency (figures 3 and 4).

**Table 2.** Symbols used and definitions

| Symbol | Concept |
| --- | --- |
| $J_{x_i, y_i}$ | Community size of patch $i$ with coordinates $x_i$ and $y_i$ |
| $\mathcal{P}$ | Total number of patches |
| $\mathfrak{d}_c = \mathfrak{d}_o + \mathcal{A} \sin(\pi \mathfrak{f} t)$ | Dispersal radius to connect two patches |
| $\hat{\mathfrak{d}}_c$ | Mean dispersal radius to connect two patches |
| $\mathfrak{d}_o$ | Initial dispersal radius |
| $\mathcal{A}$ | Amplitude of change in the dispersal radius |
| $\mathfrak{f}$ | Frequency of change in the dispersal radius |
| $\mathcal{G}$ | Generation time or complete turnover in the landscape |
| $\mathfrak{d}_{ij}$ | Geographical distance between patch $i$ and $j$ |
| $\Gamma_i = \sum_{j \neq i} (\mathfrak{d}_{ij} < \mathfrak{d}_c)$ | Connectivity patch $i$ |
| $m$ | Emigration rate |
| $\nu$ | Immigration rate from the regional species pool |
| $\lambda$ | Local birth rate |
| $\mu$ | Local natural mortality |
| $N_i^k$ | Abundance of species $k$ in patch $i$ |
| γ-richness | Number of species in the landscape |
| $\hat{\gamma}$-richness | Mean number of species in the landscape |
| $m_{ij}^k = (m/\mathfrak{d}_{ij})(N_j^k/J_{x_j, y_j})$ | Dispersal from patch $j$ to $i$ for species $k$ with abundance $N_j^k$ |
| $\mathcal{C}$ | Number of components in the landscape |
| $\hat{\mathcal{C}}$ | Mean number of components in the landscape |

**Figure 2:**
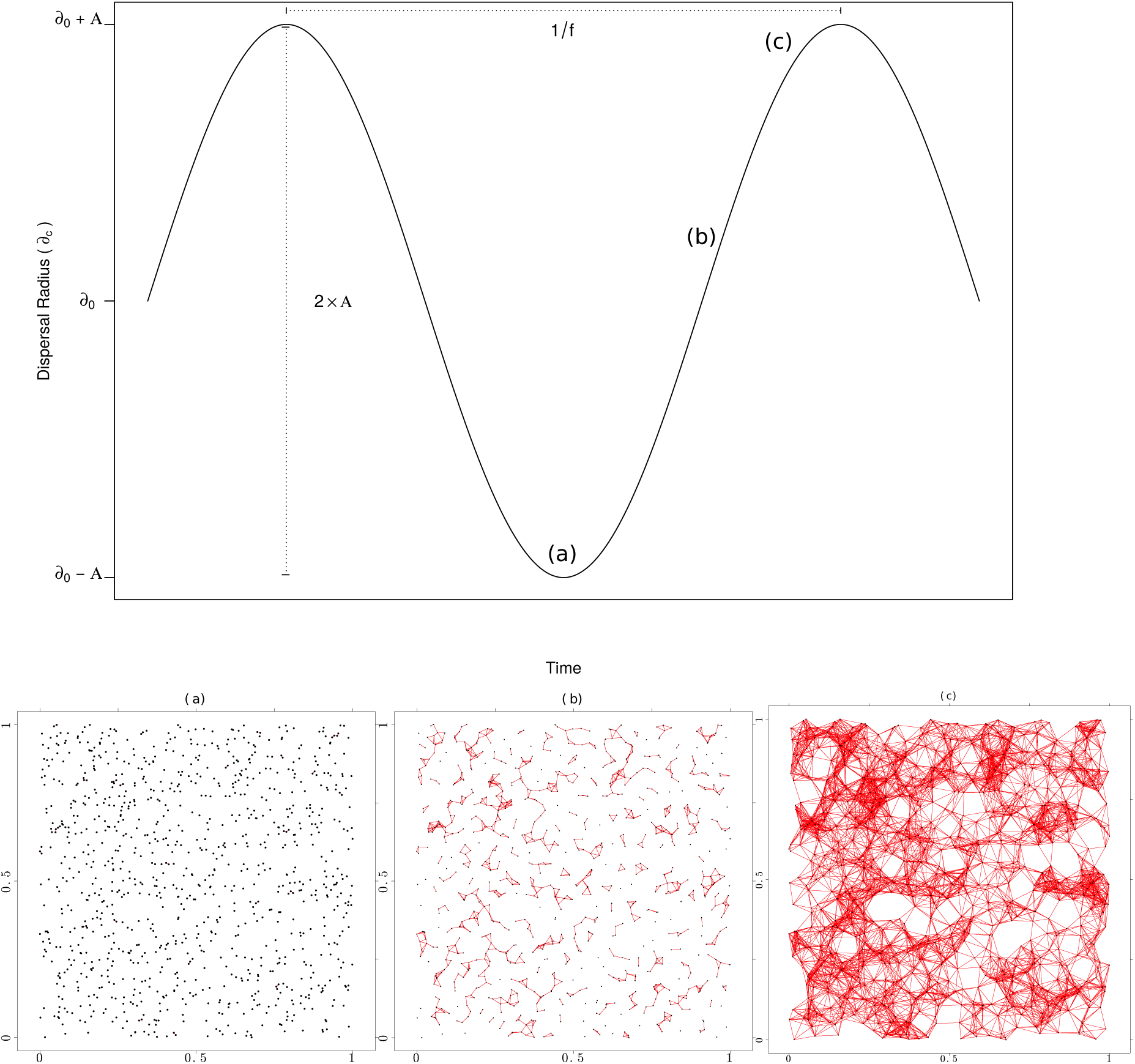
Top) Dispersal radius (*d*_c_) to determine whether two patches are connected, as a function of initial dispersal radius, *d*_0_, amplitude, *𝒜*, and frequency, f. The value of *d*_c_ fluctuates around *d*_0_, with a period given by the inverse of the frequency, f. Maximum fluctuation is given by 2*𝒜*. (a), (b), and (c) represent low, medium, and high values of dispersal radius, respectively, and are related to the topology of the networks in the bottom figure. Bottom) Landscape connectivity in three scenarios of low dispersal radius and landscape connectivity, (a), medium dispersal radius and landscape connectivity, (b), and large dispersal radius and landscape connectivity, (c). Two patches *i* and *j* are connected if their geographical distance, *d*_ij_, is lower or equal than the dispersal radius, *d*_c_.

**Figure 3:**
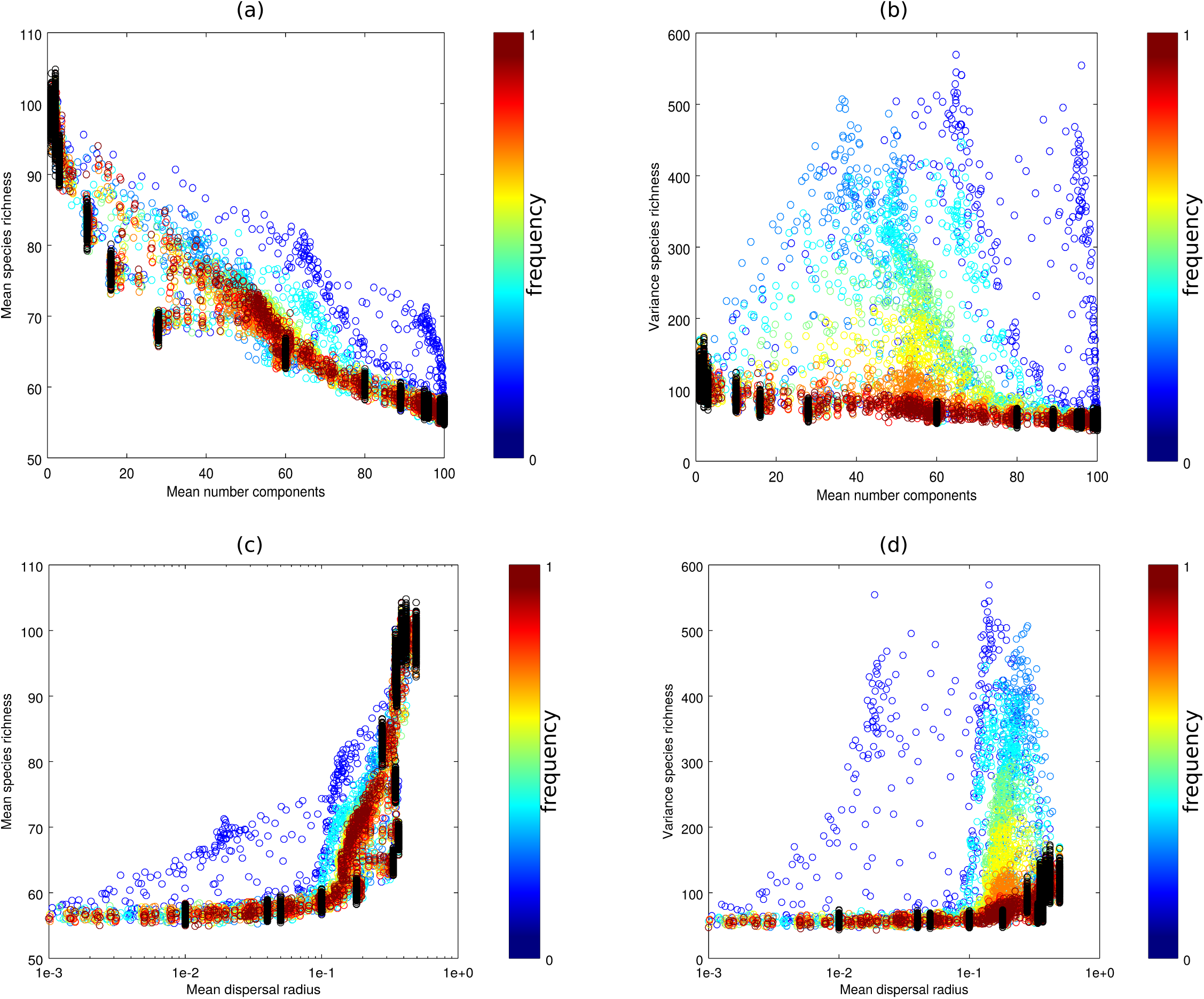
Mean (panels a and c) and variance (panels b and d) of regional species richness (*γ*–richness) as a function of the mean number of components in the landscape (panels a and b) and of the mean dispersal radius (panels c and d). The color scale represents different frequencies, f, i.e., different connectivity dynamics (see figure 2). The mean and the variance of gamma has been computed over the last 500 generations of each replicate. Simulations were done with a migration rate, *m* = 0.3 and with an immigration rate from the species regional pool, *ν* = 0.003. The total number of patches is *𝒫* = 100, the patch size is given by *J*_*x*_*i*_,*y*_*i*__ = 100 individuals and the number of generations per replicate is *𝒢* = 1000.

**Figure 4:**
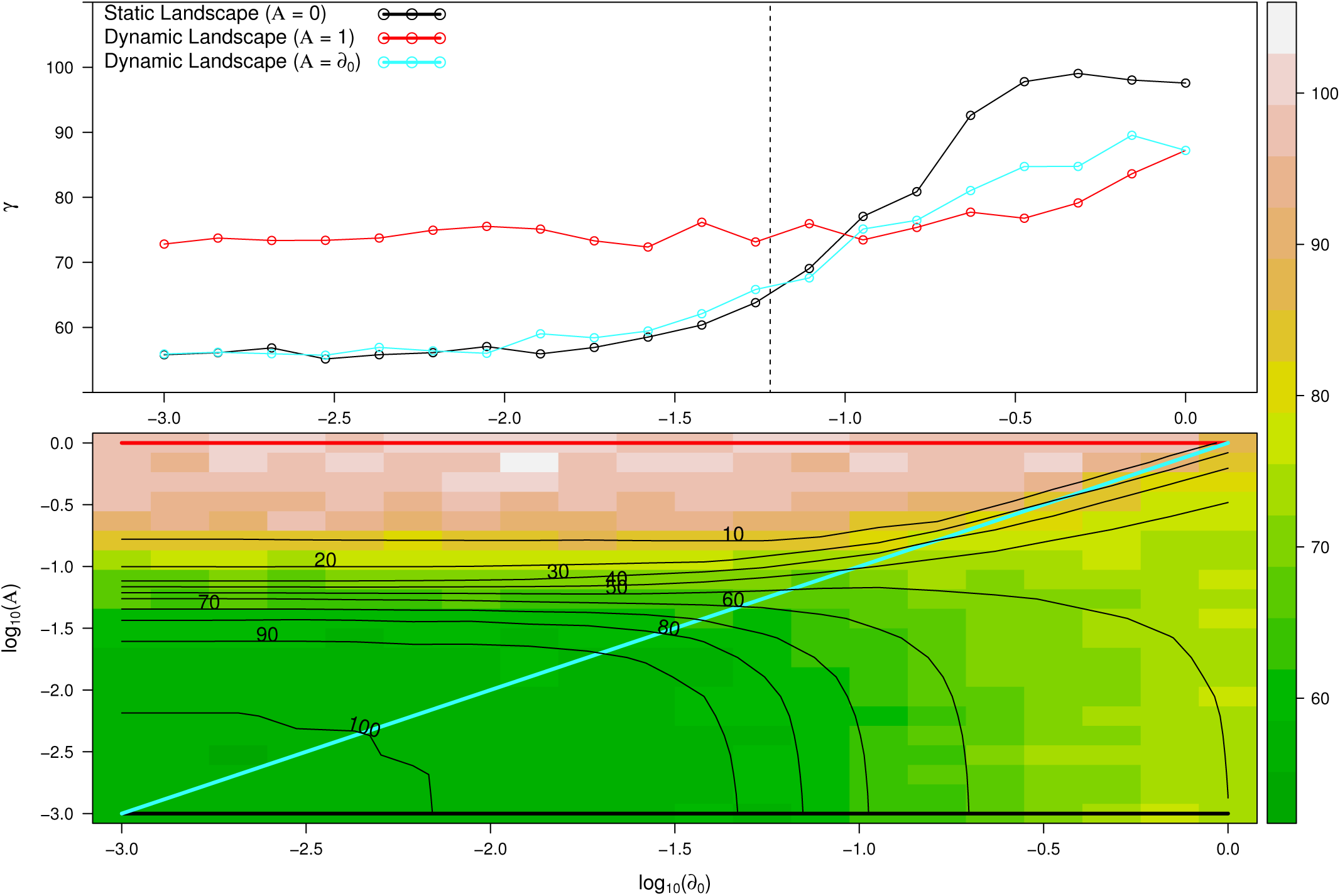
(Top) Represents *γ*–species richness as a function of the initial dispersal radius, *d*_0_, for static landscapes (black line, *𝒜* = 0), dynamic landscapes with *𝒜* = *d*_0_ (blue line), and dynamic landscapes with *𝒜* = 1 (red line). Vertical dotted line represents the critical threshold in static landscapes (see Results). (Bottom) *γ*–species richness as a function of *d*_0_ and *𝒜* for f =0.01. Isoclines (black lines) represent the mean number of components, *Ĉ*, for each combination of *d*_0_ and *𝒜*. Red, blue, and black lines represent dynamic, dynamic with *𝒜* = *d*_0_, and static landscapes, respectively, with the same values as in the top panel. Simulations were done for m = 0.3, *ν* = 0.003, *𝒫* = 100, *J*_*x*_*i*_,*y*_*i*__ = 100, *𝒢* = 1000, and f = 0.01 (see table 2).

Our results show that the number of species coexisting in fragmented landscapes differs between static and dynamic landscapes. We show that regional species richness (i.e., *γ*–species richness) decays as the landscape becomes more fragmented, both in static and dynamic landscapes, but the rate of this decay depends on the amplitude and frequency of the fluctuations of landscape connectivity (figure 3). Our results also show that for low frequency of change in landscape connectivity, the variance of regional species richness peaks with an intermediate number of isolated components in the landscape. This result suggests that high or low *γ*–species richness can occur in dynamic landscapes with a large number of components for a broad range of values of amplitude and frequency determining landscape connectivity. When varying the dispersal radius value in static landscapes, we observe a fragmentation threshold below which species richness drops dramatically (figures 3 and 4). The fragmentation threshold does not occur in dynamic landscapes for a broad range of amplitude and frequency values determining landscape connectivity (figure 4). In summary, our approach connects a mechanistic description of fluctuations of dispersal radius to landscape connectivity to explore the consequences of landscape dynamics for regional species richness.

## The model and its implementation

In this section, we describe the computational model, while the mathematical equations and further technical details are presented in the on-line supporting information (SI-B). The mathematical definitions are provided in table 2.

### Static and dynamic landscapes

We use a spatially explicit individual-based model in patchy and dynamic landscapes. We run our simulations in landscapes consisting of randomly located sites with range values between [0,1] representing landscapes of any possible scale. Each patch *i* has a spatial location given by the coordinates (*x*_*i*_, *y*_*i*_). Two patches *i* and *j* are connected by individuals dispersing if their geographic distance, *d*_ij_, is equal or smaller than a threshold distance (i.e., dispersal radius), *d*_c_. This dispersal radius is fixed in static landscapes and follows a sinusoidal signal in dynamic ones. Dispersal radius to connect patch *i* and *j* follows:

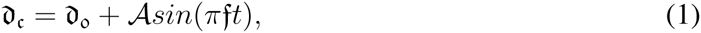

where *t* is time and *d*_0_, *𝒜*, f are the initial dispersal radius, the amplitude and the frequency of the landscape respectively. In figure 2 we show a graphical representation to visualize the effect of amplitude and frequency on the dispersal radius and landscape connectivity (i.e., the number of connections of each patch *i* with other sites in the network changes with time, see animations SI-A3 and SI-A4). In static landscapes, the connectivity of the landscape is only a function of the initial dispersal radius, *d*_0_. As the “static” landscape name suggests, the initial dispersal radius is the only value determining the threshold to connect two patches. There is no variance related to this initial dispersal radius value, and thus there is a fixed dispersal radius given by *d*_c_ = *d*_0_.

### Population dynamics and dispersal in dynamic landscapes

In our approach there can be several species in each patch and the state of each patch is described by a vector of species abundances. To model spatio-temporal changes in the abundance of these patches, we need to define dispersal rules together with population dynamics. We assume that all patches are of the same size and habitat type; we do not associate a priori a value for each patch which determines the habitat type as, for example, Rybicki & Hanski (2013) do. Instead, we allow individuals to disperse between any two patches only as a function of species abundance of the leaving patch. In this scenario individuals only can move between connected *i* and *j* patches (i.e., those patches satisfying the condition *d*_ij_ ≤ *d*_c_). At the beginning of the simulations we have an initial population that spreads instantaneously across the whole landscape. We assume that all patches are fully occupied and have the same carrying capacity, i.e., population size at a given patch *i*, *J*_*x*_*i*_,*y*_*i*__, is equal to the patch environmental carrying capacity. The total number of individuals in the landscape is *J* = *J*_*x*_1_,*y*_1__ + *J*_*x*_2_,*y*_2__ + *J*_*x*_3_,*y*_3__ + *J*_*x*_4_,*y*_4__, …, +*J*_*x*_*𝒫*_,*y*_*𝒫*__, with *𝒫* the total number of patches.

Population dynamics on the spatial network occur under a zero-sum birth and death process in overlapping generations. This means that at each time step an individual dies from a randomly chosen patch *i*. This individual is replaced with an individual coming from another patch (i.e., migrant), from the same patch than the death individual or from the regional species pool. Parents are chosen with probability *m* from outside patch *i* within the network, with probability *ν* from the regional species pool, or with probability *λ* (i.e. local birth rate), defined as *λ* = 1 − *m − ν*, from the patch *i*. We consider an extremely diverse regional species pool containing an infinite number of species. Because of the infinite number of species in the regional pool, we assume that every immigration event introduces a new species. Immigration of a new species corresponds to speciation in the context of metacommunity models (Vanpeteghem & Haegeman (2010)). Dispersal from patch *j* to patch *i* of species *k* is defined by:

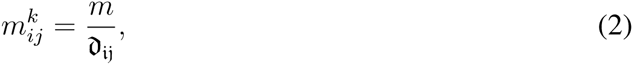

with *d*_ij_ the geographical distance between patch *i* and *j* satisfying *d*_ij_ ≤ *d*_c_ and *m* is the intensity of emigration rate. Because dispersal from patch *i* to patch *j* is the same as in the opposite direction 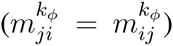, this represents symmetric, patch- and density-independent dispersal where dispersal to connected and less distant patches is more likely than dispersal to more distant patches.

### Implementation and simulations

Prior to the simulations, one needs to specify the parameters for generating the landscape and the regional pool of species. The landscape is generated following a 2D-random geometric network as described in the section “Static and dynamic landscapes”. Simulations were carried out with an initial population at each patch *i*, *J*_*x*_*i*_,*y*_*i*__, of 100 individuals for a total of 100 patches. The population size and the number of patches remained constant throughout the simulations. Results for figures 3-4, and SI-C1 to SI-E1 were obtained after 100 replicates with 1000 generations each, where a generation, *𝒢*, is an update of the total number of individuals, *J*, in the landscape. Values plotted represent the mean and the variance across the last 500 generations per replicate. We explored a broad range of parameter values from a uniform distribution with values *𝒰* [0.001, 1] for the initial dispersal radius, *d*_0_, the amplitude, *𝒜*, and the frequency, f. We set mortality rates equal to 1 (i.e., the natural mortality rate, *μ*). Rates of immigration from the regional species pool, *ν*, and the intensity of emigration rate, *m*, were set to 0.003, and 0.1, respectively. Local birth rates for each metacommunity, *λ* = 1 *− ν − m*, so that a new individual replacing the dead individual appears with certainty.

### Landscape connectivity and *γ*–species richness

We calculated the mean number of components per replicate as a proxy of landscape connectivity and availability together with the mean and variance regional species richness (i.e., *γ*–species richness) for the simulations with static and dynamic landscapes (figure 3 and SI-C1). We remark that a component can be formed by one or several isolated patches (table 1). We also calculated mean and variance of the *γ*–species richness as a function of the dispersal radius, *d*_c_ (figure 3 and SI-D1). We also performed spectral analysis on the time series of *γ*–species richness, in order to detect possible resonance between fluctuation of the landscape and species richness. We plotted the mean and variance *γ*–species richness and also the mean number of components vs. all amplitudes, *𝒜*, frequencies, f, and initial dispersal radius, *d*_0_, explored (figure 4 and SI-E1).

## Results

We found that migration, and the frequency and amplitude of the dispersal radius play a key role in predicting regional species richness. For medium to high migration rates, *m* = 0.3, the mean regional species richness decayed with the increasing mean number of isolated components in static landscapes (figure 3 top left, black circles). The overall trend for the mean regional species richness for dynamic landscapes was qualitatively similar to static landscapes with the mean regional species richness decaying with an increasing number of isolated components in the landscape for all values of frequency (figure 3 top left, Spearman-*ρ* > 0.37, All p < 0.05). However, mean regional species richness values differed between high and low frequency (compare high frequency in red, orange and yellow, f *>* 0.1, with low frequency, f = [0.001,0.1], light blue and dark blue in figure 3). For example, static and dynamic landscapes with high values of frequency (frequencies in red, orange and yellow, f *>* 0.1) predicted less than 70 species in a highly fragmented landscape containing 60 components. Predictions of the mean regional species richness for dynamic landscapes with low frequency values, f = [0.001,0.1] reached values above 80 species (light and dark blue in figure 3 top left). Landscapes with frequency values equal to 0 recover a static landscape (equation 1) and the dispersal radius, *d*_c_, is equal to the initial dispersal radius, *d*_0_, as for static landscapes. The trend of decreasing regional species richness with the number of components in the landscape observed for high migration rate was less strong with low and very low migration rate values (*m* = 0.1 and *m* = 0.01, respectively, figure SI-C1 top and bottom left).

The variance for static landscapes followed the same pattern as the mean regional species richness with the number of components for high migration rates and high values of frequency (*m* = 0.3, figure 3 top right, black circles and red, orange and yellow circles, f *>* 0.1, Spearman-*ρ* > 0.31, All p < 0.05). However, the variance for dynamic landscapes peaked and showed no correlation approximately at an intermediate number of isolated components in the landscape for medium and low frequency values of change in landscape connectivity (figure 3 top right, f = [0.001,0.1], represented as light green, light blue and dark blue, Spearman-*ρ* < 0.18, p > 0.1). This result suggests that high or low *γ*–species richness occurred in dynamic landscapes with a large number of components and for a broad range of values of amplitude and frequency determining landscape connectivity. The decay of the variance of regional species richness with the number of components in the landscape for low migration dynamics reproduced the pattern observed for the mean species richness for static landscapes (figure SI-E1 top and bottom right). Our results showed that an increasing number of fragments in the landscape predicted less regional species richness. However, we show also that the decay in the mean and variance of regional species richness is affected by both differences in migration rates and by differences in the frequency of change of the dispersal radius. Thus, migration and connectivity dynamics played a key role to predict the regional species richness in dynamic landscapes with fluctuations in landscape connectivity supporting metacommunities with higher mean and variance in species richness than the observed richness in static landscapes.

Our analysis of the relationship between dispersal radius and *γ*–species richness showed a fast decay in species richness for high migration rates in static landscapes (figure 3 bottom left, black circles). The threshold observed in static landscapes decayed less strongly in dynamic landscapes for low frequency and low migration rate values (figures 3 bottom left, frequency values, f = [0.001,0.1], represented as light green, light blue and dark blue and figure SI-D1 top left for migration rate, *m* = 0.1). The threshold in species richness was not observed for very low migration rates (figure SI-D1 bottom left for migration rate, *m* = 0.01). For low migration rate values, both static and dynamic landscapes show the same uncorrelated pattern with the mean and variance regional species richness varying greatly for the range of dispersal values explored (Spearman-*ρ* < 0.15, p > 0.1). High variance in regional species richness was observed in dynamic landscapes with low values of frequency, f = [0.001, 0.1], for large mean dispersal values (figure 3 bottom right, light green, light blue and dark blue). Moreover, performing spectral analysis on the time series of *γ*–species richness, we found no particular correlation between the frequency of the landscape and the characteristic frequencies of species richness. Our results showed that medium to high migration rates predicted stronger deviations from static landscapes and faster decay of species richness and overall a lower species richness when decreasing the dispersal radius than low or very low migration dynamics (compare figure 3 with SI-C1 and SI-D1).

To explore the robustness of the decay of regional species richness with the number of components in the landscape in static and dynamic landscapes we simulated a broad range of amplitude, *𝒜*, frequency, *f*, and initial dispersal radius values, *d*_0_ (figure 4 and SI-E1). The rapid decay of landscape connectivity with decreasing dispersal radius followed from the predicted analytical percolation threshold in random geometric graphs. The critical threshold in our landscape is given by *D*_*c*_ = *L ×* sqrt(4.52/(4 *× π × 𝒫*)) = 0.06 (figure 4, vertical dotted line, log10(0.06) = -1.22), where *L* is 1, and *𝒫* the number of patches, 100. Below this critical threshold, the landscape was fragmented into a large number of disconnected components and *γ*–species richness was more strongly reduced in static and dynamic landscapes for a broad range of frequency values (figure 4 and SI-E1 top compare static landscapes, black line, with dynamic landscapes, red lines; blue line shows dynamic landscapes with *𝒜* = *d*_0_). These results were robust to changes in the frequency determining the dispersal radius (figure SI-E1 for frequency, f = 0.001 (a), 0.01 (b), 0.1 (c) and 1 (d)). The threshold decreasing *γ*–species richness in static landscapes did not occur in dynamic landscapes. This result remained qualitatively similar for two orders of magnitude of frequency values (figure SI-E1 for frequency, f = 0.001 (a), 0.01 (b), 0.1 (c)). In summary, the fast decay in species richness as the landscape becomes fragmented in static landscapes did not occur in dynamic landscapes for a broad range of amplitudes, *𝒜*, frequencies, f, and initial dispersal radius values, *d*_0_. This suggests that dynamic landscapes may support metacommunities with higher species richness than static landscapes in fragmented landscapes.

## Discussion

Our study adds to previous attempts to connect species persistence to dynamic landscapes (Hanski (1999); Keymer *et al.* (2000)). Among the many factors driving landscape connectivity we focus on the periodic ones. Different periodicity can be described by varying the amplitude and the frequency of the change in landscape connectivity. Here we described how the amplitude and the frequency of landscape connectivity drive coexistence in multispecies communities. Our results show that the fluctuations of landscape connectivity support metacommunities with higher species richness than static landscapes (figures 3-4). We show the decay in the mean and the variance of regional species richness, caused by increasing number of fragments, strongly differed between low, medium and high migration rates and between different values of the frequency values driving landscape connectivity (figure 3). This means that highly fragmented landscapes can support a species rich metacommunity if the landscape becomes periodically connected. The positive effect of these periods of high landscape connectivity which allows dispersal and range expansions on *γ*–species richness thus offsets the negative effects of periods of low connectivity. Our results also suggest that landscapes characterized by fast changes of connectivity relative to the generation time of organisms predict qualitatively the same outcomes as static landscapes (i.e., landscape with high frequency, figure 3). This result implies that analytical predictions obtained from the classical metacommunity theory in static landscapes may be valid for rapidly changing dynamic landscapes with high frequencies determining dispersal dynamics of populations (figures 3 and 4). However, we have also shown that there is a broad range of frequency and amplitude values which provide predictions that strongly differ from static landscapes.

Contrary to our metacommunity model, classical studies of predator-prey and competitive interactions reported that higher landscape connectivity and migration rates tend to homogenize metacommunities and decrease species richness (Ellner *et al.* (2001); Fox *et al.* (2011)). High landscape connectivity in predator-prey systems tends to destabilize prey populations, which leads to extinctions and thus decreases species richness (Ellner *et al.* (2001); Fox *et al.* (2011)). Similarly, competitive communities with highly connected landscapes tend to have only a few dominant species (Holyoak *et al.* (2005)). These results follow from interaction asymmetries, which are not included in our models. Instead, the models we have explored here emphasize random and limited dispersal and demographic stochasticity, as the main drivers of metacommunities in dynamic landscapes. Our approach did not explicitly test for directionality of migration or selection and we assumed equal growth rates across the landscape, nor did we assume any asymmetry in competition or trophic interactions as possible mechanisms for structuring diversity in our static and dynamic landscapes, hence a neutral theory of biodiversity in dynamic landscapes was applied. While our model assumes neutral dynamics and random geometric graphs for population and migration dynamics, in a more realistic scenario we expect more differences between static and dynamic landscapes. For example, in our model all the individuals and species use the available connections between patches equally, but niche differences within and between species, different habitat preferences or landscape heterogeneity may provide a more strict threshold for the decay of *γ*–species richness. Our prediction of high regional species richness in landscapes with patches alternately isolated and then highly connected for periods of time is tentatively supported by studies of river systems which show that even brief periods of increased connectivity may lead to gene flow with significant effects on genotypic diversity of populations over the landscape (Boizard *et al.* (2009)). In the metacommunity context, brief periods of high landscape connectivity may allow local species to spread rapidly to a number of new sites providing opportunities for population growth and rescue from extinction by demographic and environmental stochasticity in small local populations.

Extinction thresholds form one of the core predictions from metapopulation and metacommunity theory (Tilman *et al.* (1994); Bascompte & Solé (1996); Keymer *et al.* (2000); Fahrig (2002); Ovaskainen & Hanski (2003); Rybicki & Hanski (2013)). Several models and field data have shown single and multiple species extinction thresholds with increasing habitat loss in random and nonrandom habitat destruction scenarios (Fortuna & Bascompte (2006)). While most of the studies dealing with habitat destruction change the total amount of available habitat, the extinction threshold obtained in our approach is produced in landscapes with constant total amount of available landscape. Despite this difference in the approach used to understand regional species richness with increasing landscape fragmentation, our result show that the classical percolation threshold found in random geometric landscapes predicts a multiple species extinction threshold in static landscapes. However, this percolation threshold does not predict a multiple species extinction threshold in dynamic landscapes (figure 4). This means dynamic landscapes allows for dispersal periods that compensate for local extinctions during periods of low connectivity.

Microcosm or mesocosms experiments with contrasting regimes of amplitude and frequency determining connectivity fluctuations could be used to test our predictions under laboratory conditions. Model systems like bacteria, protists (Carrara *et al.* (2012); Altermatt *et al.* (2015)), small invertebrates such as zooplankton (Steiner *et al.* (2011)) or insects (Govindan & Swihart (2012)) may provide a good level of control over the landscape-level parameters to test predictions from dynamic landscapes models. Long-term field data can also be used to explore landscape dynamics models incorporating more realistic climatic regimes or broader geographic regions in deep time to infer the amplitude and frequency (or additional parameters capturing fluctuations at different temporal scales) that best predict the spatio-temporal fluctuations in species diversity. For example, there is evidence of rapidly changing landscapes in the Arctic and Antarctic regions with the ice cover dynamics (animations SI-A1 and SI-A2), but the amplitude and frequency required to predict such fluctuations and their impact on local and regional species richness are currently unknown. Landscape dynamics approximations can also help to discern how much complexity is required to make predictions that fit periods of peaks or flattened species richness gradients as observed in the fossil record for some periods of the latitudinal biodiversity gradient (Mannion *et al.* (2014)). In deep time, transitions between habitat types at the continental scale occurring during glacial-interglacial cycles over long temporal scales would require to include non-periodic landscape dynamics (i.e., plate tectonic or continental drift) (Werneck *et al.* (2011)) and here we provide a simple model that can be extended to include those more realistic scenarios.

### Future perspectives

Given the rapid changes observed in natural and human-disturbed landscapes, there is a growing need to develop methods that more accurately describe the effects of dynamic landscapes in metacommunities. Here we have developed an individual-based metacommunity model to explore the effect of amplitude and frequency of fluctuations of organisms’ dispersal radius on local and regional species richness. In addition to temporal fluctuations of dispersal radius (equation 1 and equations in SI-B), we can simulate destruction of patches and creation of new patches at random (or seasonal) time points. Similarly, spatial heterogeneity or temporal fluctuations in the carrying capacity of individual patches could also be included. We can thus start to explore the interactive effects of patch and connectivity dynamics on local and regional species richness. In the absence of patch dynamics, our results show that the fluctuations of landscape connectivity support metacommunities with higher species richness than static landscapes in fragmented landscapes but the combined effect of patch and connectivity dynamics can change these predictions. Future research would need to combine patch and connectivity dynamics to further advance our understanding of short- and large-scale patterns of biodiversity changes in rapidly changing landscapes.

## Acknowledgments

This study was supported by the Swiss National Science Foundation project 31003A-144162 (to CNdeS and CM), by the Sciex fellowship project 12.327 (to JK) and by the Swiss National Science Foundation International Short Visits project IZK0Z3_158668 (to CM and JK). JK is also supported by the Czech Science Foundation (project GP14-10035P).

### Appendix A from Charles Novaes de Santana, J. Klecka, G. M. Palamara, and C. J. Melián, Metacommunities in dynamic landscapes

#### Landscape dynamics animations

##### SI-A1; Sea Arctic ice cover animation (ArcticSI-A1.avi)

This animation shows monthly Arctic sea ice cover for the period between October-1979 to September-2010, downloaded from Cavalieri *et al.* (1996)

##### SI-A2; Sea Antarctic ice cover animation (AntarcticSI-A2.avi)

This animation shows monthly Antarctic sea ice cover for the period between October-1979 to September-2010, downloaded from Cavalieri *et al.* (1996)

##### SI-A3; Dynamic landscape animation (figureRGNSI-A3.avi)

This animation shows fluctuations in landscape connectivity using amplitude, *𝒜*, frequency, f, and initial dispersal radius, *d*_o_, of 0.15, 0.2 and 0.15, respectively (left, scenario 1 with *𝒜* ≠ *d*_o_), and 0.3, 0.2, and 0.15, respectively (right, scenario 2 with *𝒜* ≠ *d*_o_).

##### SI-A4; Dynamic landscape animation (figureRGNSI-A4.avi)

This animation shows fluctuations in landscape connectivity using amplitude, *𝒜*, frequency, f, and initial dispersal radius, *d*_o_, of 0.4, 0.05 and 0.4, respectively (left) and 0.4, 0.25, and 0.4 (right), respectively.

### Appendix B from Charles Novaes de Santana, J. Klecka, G. M. Palamara, and C. J. Melián, Metacommunities in dynamic landscapes

#### Stochastic metacommunity landscape dynamics

Here, we explain in detail how we combine dispersal with local population dynamics. The following equations conceptualize metacommunity dynamics. The first (second) equation gives the transition probability that the *k*^*th*^ species of metacommunity declines (increases) in abundance by one individual in patch *i*

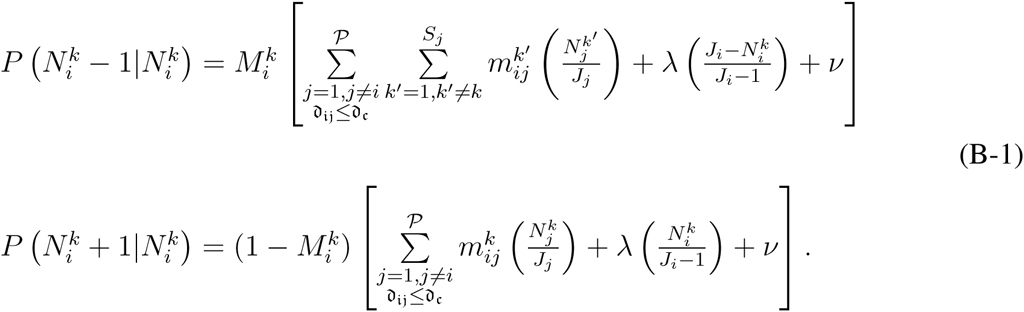

Here 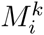 describes density-dependent mortality rate of species *k* in patch *i*. This mortality is the natural per capita mortality rate described in this article by 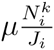. 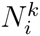 and *J*_*i*_ are the total number of individuals of species *k* in patch *i* and the total number of individuals in patch *i*, respectively. *𝒮*_*j*_ and *𝒫* are the total number of species in patch *j* and the total number of patches, respectively. In addition to the mortality rate parameters, there are three more metacommunity specific parameters: *λ*, the local birth rate, *m*, the intensity of emigration rate, and *ν*, the immigration rate from the regional species pool.

The first equation in (B-1) gives the transition probability for the *k*^*th*^ species to decline in abundance by one individual in patch *i*. For this to happen, an individual must die in the *k*^*th*^ species, which occurs at a rate given by 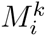. The first probability inside the brackets is that of an immigration event of some species other than *k* from a patch different to *i* (see equation 2 in the main text with *d*_ij_ the geographical distance between patch *i* and *j* satisfying *d*_ij_ ≤ *d*_c_). The second term represents the probability of having a local birth in a species other than *k* with the -1 subtracted in the denominator after the death in the previous step of one individual in this patch. The third term describes the probability of an immigration event from the regional species pool. The second equation in (B-1) describes the transition probability for the *k*^*th*^ species to increase by one individual. For this to happen, there must be no local death in species *k* which is given by 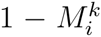. The other terms in brackets stand for dispersal (the first term), local birth of an individual of species *k* (second term), and immigration of a new species *k* from the regional species pool. This last event can occur only when there was no such species, i.e., when 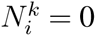 at time *t* #x2212; 1.

### Appendix C from Charles Novaes de Santana, J. Klecka, G. M. Palamara, and C. J. Melián, Metacommunities in dynamic landscapes

#### Mean and variance *γ*–species richness as a function of the mean number of components

**Figure C1.**
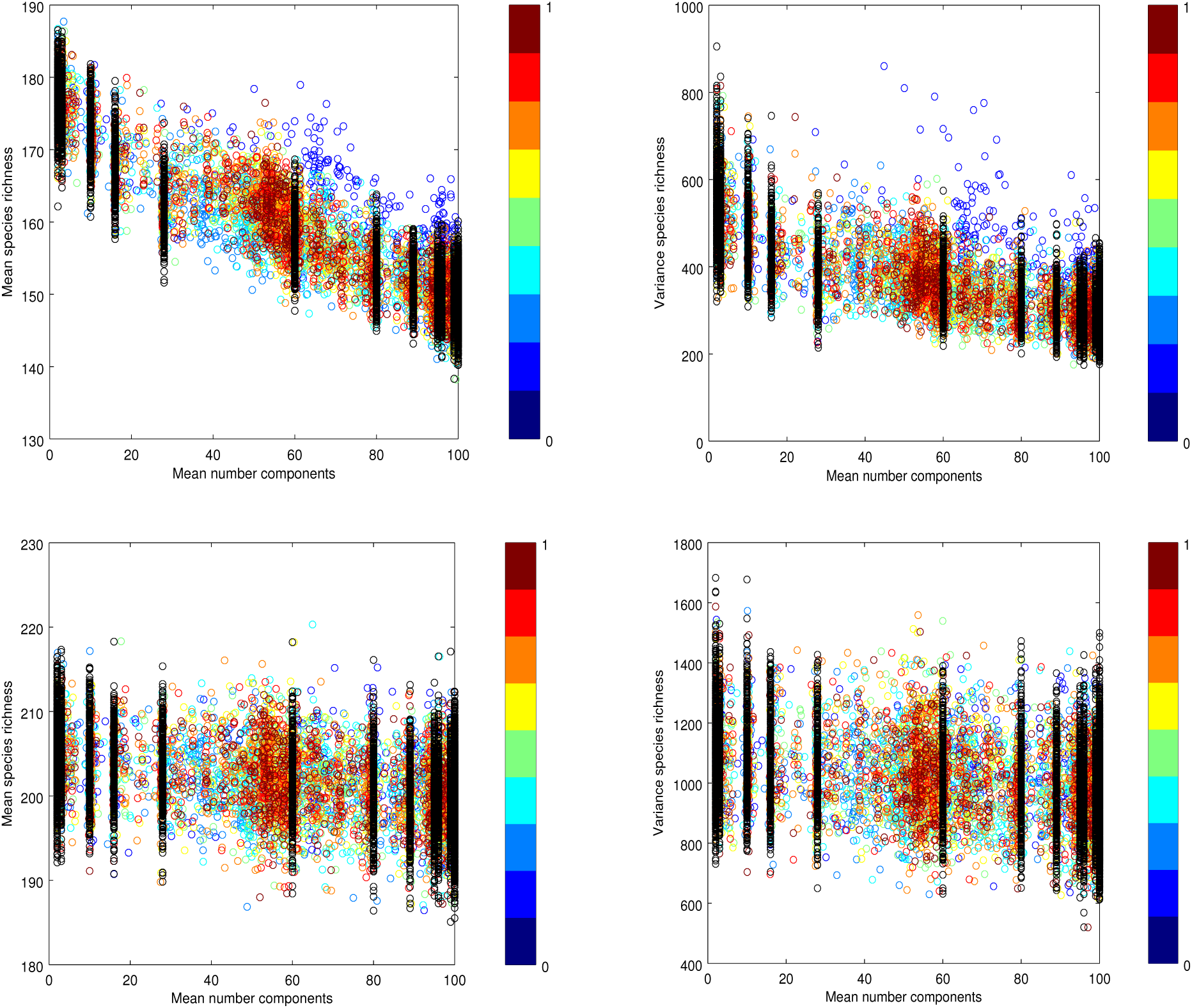
This figure shows the mean (left) and variance (right) *γ*–species richness as a function of the mean number of components for two migration rates, m = 0.1 (top) and m = 0.01 (bottom). Parameter values used are as in figure 4 of the main ms.

### Appendix D from Charles Novaes de Santana, J. Klecka, G. M. Palamara, and C. J. Melián, Metacommunities in dynamic landscapes

#### Mean and variance *γ*–species richness as a function of mean dispersal radius

**Figure D1.**
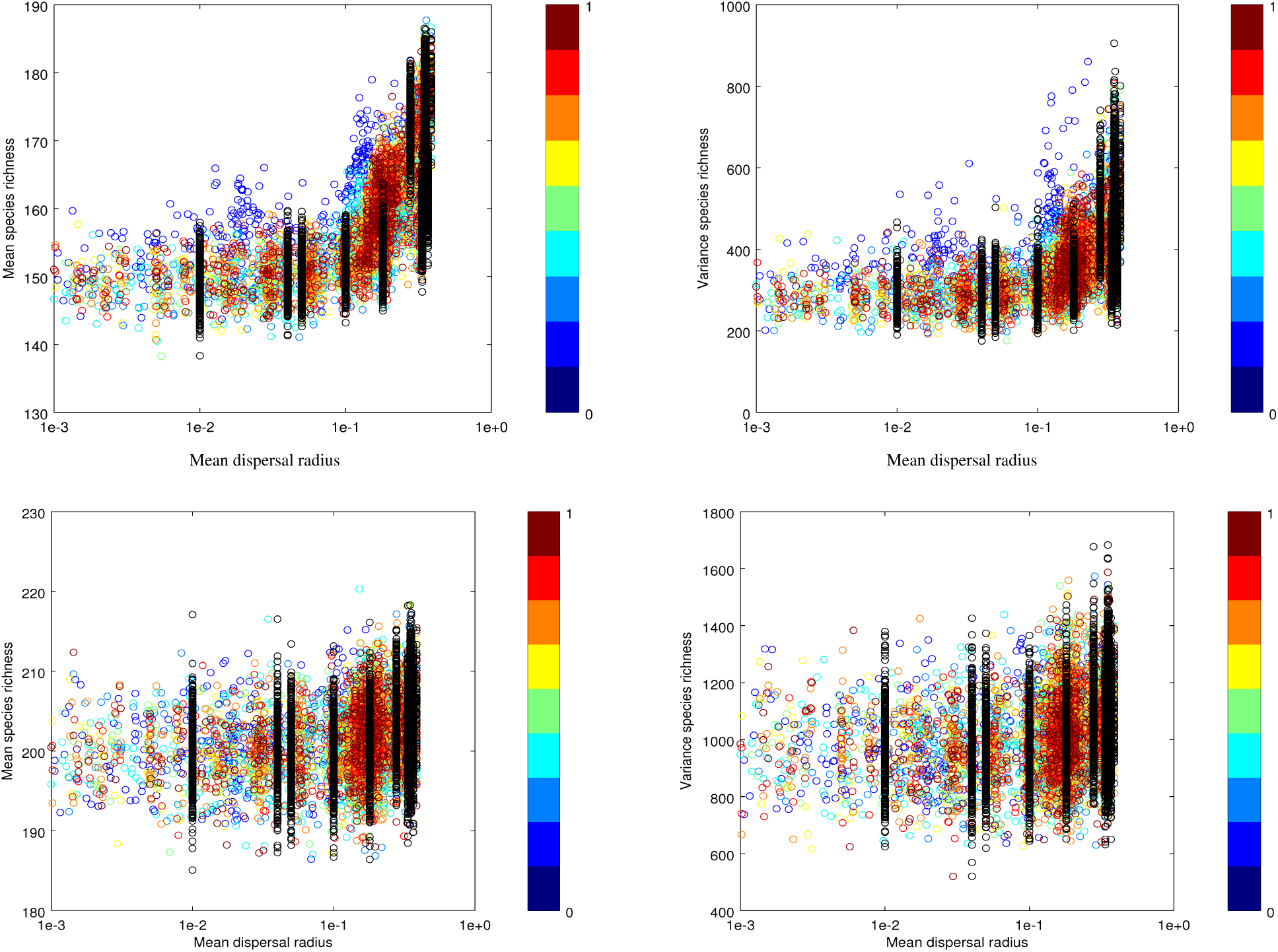
This figure shows the mean (left) and variance (right) *γ*–species richness as a function of the dispersal radius, *d*_c_, for two migration rates, m = 0.1 (top) and m = 0.01 (bottom). Parameter values used are as in figure 5 of the main ms.

### Appendix E from Charles Novaes de Santana, J. Klecka, G. M. Palamara, and C. J. Melián, Metacommunities in dynamic landscapes

#### Mean *γ*–species richness

**Figure E1.**
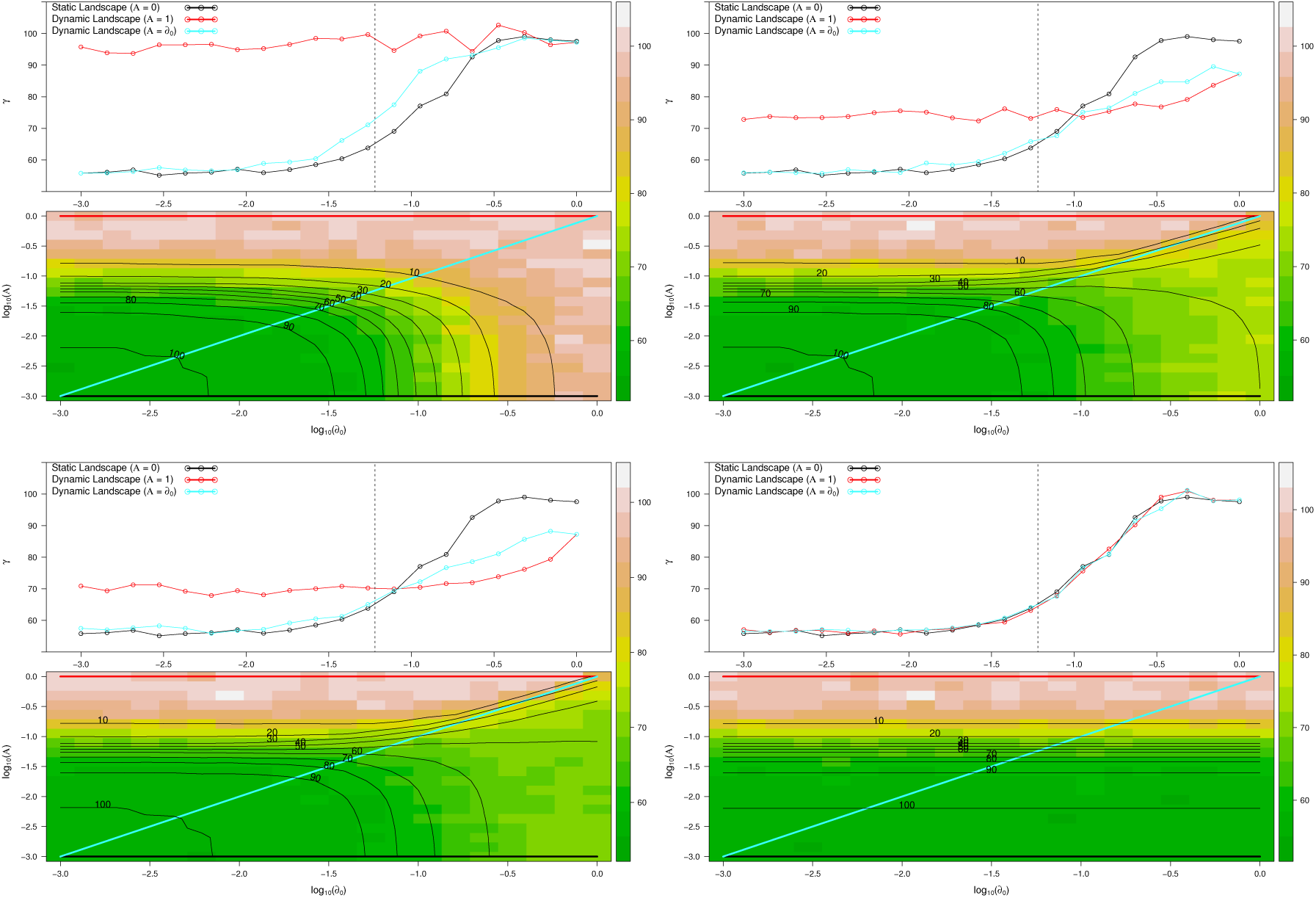
This figure shows the mean *γ*– species richness as a function of the dispersal radius, *d*_c_), and amplitude, *𝒜*, for static landscapes (black line, *𝒜* = 0), dynamic landscapes with *𝒜* = *d*_0_ (blue line), and dynamic landscapes with *𝒜* = 1 (red line) for four frequency, f, values: (a) 0.001, (b) 0.01, (c) 0.1, and (d) 1. Vertical dotted line represents the critical threshold in static landscapes. Isoclines (dotted lines) represent the mean number of components, ĉ, for each combination of dispersal radius, *d*_c_, and amplitude ν. Simulations were done for emigration rate, m = 0.1, immigration rate from the species regional pool, *ν* = 0.003, total number of patches, 𝒫 = 100, patch size, *J*_*x*_*i*_,*y*_*i*__ *ρ* = 100 individuals and number of generations per replicate, 𝒢 = 1000. Values plotted represent the mean values over the last 500 generations in each replicate.

